# Disentangling the Effects of Weather and Density on the Dynamics of a Fungal Pathogen

**DOI:** 10.1101/088054

**Authors:** Colin Kyle, Eli Goldwyn, Greg Dwyer

## Abstract

Nonlinear fitting algorithms have illuminated the role of weather in human diseases, by allowing for robust tests of mechanistic transmission models, but a lack of data has prevented applications to animal diseases. This is important because classical models that neglect weather predict that there will be a host density threshold, below which epidemic intensity will be slight, but models that include weather predict that this threshold will often be obliterated by weather variability. To test the applicability of thresholds to animal diseases, we estimated infection rates of the fungal pathogen *Entomophaga maimaiga* in the gypsy moth, by collecting larvae during epidemics at a range of host densities and weather conditions, and we estimated the pathogen's force of infection, by exposing experimental larvae to the pathogen for 24 h periods in the field. By fitting a range of models to our data, we show that epidemics of this pathogen are best explained by a model that allows for positive effects of both host density and cool, moist weather on transmission, such that weather-only and density-dependence-only models provide vastly poorer explanations for the data. Despite the effects of weather, the combined model shows that the effects of density in *E. maimaiga* are strong enough to ensure that the density threshold will have important effects on the probability of epidemics. Our work shows that weather and density-dependent transmission can interact in non-intuitive ways, and provides an illustration of the usefulness of nonlinear fitting for understanding animal diseases.

---

Efforts to understand the effects of weather on human infectious diseases have seen significant advances through the application of nonlinear model fitting and high-performance computing [1]. Nonlinear fitting, however, has seen few applications to animal diseases, even though statistical associations between weather and the spread of animal diseases have been widely documented [2]. Our understanding of the effects of weather on animal diseases is therefore very limited. Part of the problem is that most data sets on animal diseases are insufficient to allow estimation of the parameters of mechanistic models, especially for the vertebrate diseases that are the focus of most research [3]. In the few cases for which the requisite data *are* available, the models fit to the data have typically allowed for demographic stochasticity, the randomizing effects of small population sizes, but not the environmental stochasticity that arises from variable weather conditions [4].

Previous work on weather and animal diseases has therefore relied on correlational approaches based on linear or generalized linear statistical models [2]. Linear models are computationally convenient, but do not allow for explicit mechanisms. In particular, linear models inherently assume that transmission is density-independent, an assumption that eliminates the disease-density thresholds that are a basic prediction of mechanistic models [5]. Because animal densities in nature are often highly variable [6], disease-density thresholds could play a key role in determining whether epizootics (i.e., epidemics in animals) occur, but only if the effects of density are not overwhelmed by the effects of weather. Efforts to detect disease-density thresholds in nature, however, have again considered only the effects of demographic stochasticity [7]. Our understanding of the importance of thresholds in animal host-pathogen systems is therefore very limited.

Models that include the combined effects of weather and density-dependence show a wider range of behaviors than either weather-only or density-dependence only models [8], but have not been subject to robust tests with data for animal pathogens. For combined models, the problem of a lack of data is particularly acute, because robust model tests require data over a range of both host densities and weather conditions [9]. Moreover, most data sets include only observational data, but inferring mechanisms from observational data alone can be very difficult, even for diseases for which extensive data sets are available [1]. For insect pathogens in contrast, epizootics often occur at easily observable scales [10], while the small size of the host makes experimentation straightforward [11]. For fungal insect pathogens in particular, variability in host density is often paired with strong effects of weather [12]. Insect-fungus interactions are thus useful for studying how weather variability modulates the effect of host density on disease dynamics.

We therefore collected data on epizootics of the fungal pathogen *Entomophaga maimaiga*, which infects the gypsy moth, *Lymantria dispar*. The gypsy moth is an important introduced pest of hardwood forests in North America [13], but the introduction of *E. maimaiga* [14], in combination with he earlier introduction of a baculovirus pathogen [15], has reduced defoliation. The costs of defoliation are nevertheless sufficiently high that control efforts are often intensive, with costs ranging into the millions of US dollars each year [13] A mechanistic understanding of the effects of weather and density may therefore be useful for gypsy moth pest management, by providing the ability to predict when outbreak populations will collapse without intervention, and by determining the extent to which collapses are weather-dependent.

## Significance Statement

An important prediction of classical models of disease epidemics is that there will be a host density threshold, marking the lowest host density at which epidemics occur. Alternative models that incorporate weather effects suggest that weather variability can eliminate the threshold in animal diseases, but data sets sufficient to test the models have been lacking. We collected an extensive data set for a weather-affected fungal pathogen, to test if weather eliminates the disease density threshold. Our models show that weather has strong effects on the transmission of this disease, but that there is nevertheless a meaningful disease-density threshold. Our work shows that mechanistic models can be useful for understanding animal diseases, even if weather strongly alters disease dynamics.

As is often the case with fungal pathogens [12], there is good evidence that cool, moist weather enhances the infectivity and survival *E. maimaiga* spores [14, 16]. Evidence for effects of host density, however, is conflicting, probably because the pathogen’s high dispersal rate weakens local density effects [17]. Possibly for that reason, two previous studies showed opposite effects, such that density effects *were* detected across populations separated by hundreds of kilometers [18], but were *not* detected in populations separated by less than 50km [19]. In testing for combined effects of weather and density, we therefore used populations in the state of Michigan, USA that were separated by 85-350 km along a transect that spanned more than three degrees of latitude (Supplementary Information), and we collected data for three consecutive years. In each population, we quantified host densities at the beginning of each larval season, and we recorded temperature, rainfall, humidity, and *E. maimaiga* infection rates throughout the season.

In previous studies, samples sizes were small enough that it was only possible to analyze cumulative infection rates, preventing analyses of variation in infection rates during epizootics [18, 19]. This is important because in models that include density-dependent transmission, infection rates first rise and then fall as the epizootic proceeds [6], a pattern that by definition cannot be detected in cumulative infection rate data. To detect variation in infection rates during epizootics, we therefore collected 100 larvae in each population in each week of the larval season, rearing the larvae in the lab to determine their infection status.

We also carried out an experiment in which we exposed uninfected, lab-reared larvae to the environment for a 24 hour period each week, thereby producing a weekly estimate of the pathogen’s force of infection [6], the rate at which larvae become infected. Because the experiments were short enough that none of the larvae died during the exposure period, the experimental data include only the effects of transmission, whereas the epizootic data as well include the effects of other processes. Collecting both experimental and observational data therefore improved our ability to distinguish between different mechanisms affecting the dynamics of the pathogen.

We then used our data to choose between alternative models of *E. maimaiga* dynamics, such that (1) the first model assumed that transmission depends only on density, (2) the second assumed that transmission depends only on weather, and (3) the third assumed that transmission depends on both density and weather. This approach allowed us to test for effects of density and weather, but it also produced parameter estimates for each model. We then used the parameter estimates in the models to predict epizootic intensities across a range of densities, in order to understand the extent to which weather affects disease-density thresholds in this disease.

Like many outbreaking insects [20], the gypsy moth has only one generation per year, and the disease only infects larvae, which in turn are present in our study area from approximately 1 May to 15 July. We can therefore model *E. maimaiga* epizootics using a standard, single-epizootic SEIR model, modified to include the two infectious stages of *E. maimaiga*. The first of these stages consists of an over-wintering form known as a “resting spore” [21]. Resting spore germination in the spring introduces the disease into the larval population, through the ejection of short-lived infectious spores that lead to new infections [14]. Transmission from resting spores ends in approximately mid-May, but larvae infected by resting spores go on to release conidia into the air column until near the end of the larval period, causing additional infections.

Our SEIR model therefore includes separate transmission terms for conidia and resting spores:

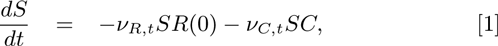

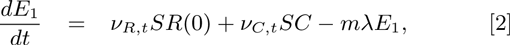

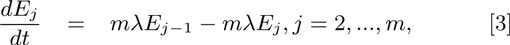

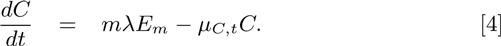

Here *S* is the density of uninfected or “susceptible” larvae per square meter, for which we calculated initial values *S*(0) using standard methods of surveying for gypsy moth densities (Supplementary Information). An epizootic in the model is then initiated by resting spores, with density *R*(0)_*i*_ in population i, which for simplicity we assume is constant for the few weeks during which resting spores bloom [18]. Larvae infected by resting spores produce conidia *C*, which lead to multiple rounds of transmission. Infected larvae transition through *m* exposed classes *E_j_* at rate λ, so that the distribution of times between infection and death is gamma distributed with mean 1/λ and variance 1/*m*λ^2^ [6]. In practice, variation in the speed of kill is low enough that we set *m =* 100 to simplify our model-fitting procedures.

In equations 1-4, the resting spore transmission rate *ν_R,t_*, the conidial transmission rate *ν_C,t_*, and the decay rate *μ_t_* are shown as functions of time, to emphasize that each may depend on variation in the weather, and on a time varying stochasticity term (in practice we considered multiple weather drivers, see Methods for the specific dependence of each parameter on the weather variables). To produce a model with density-dependent transmission but no weather effects, we assumed that each parameter varies only due to stochasticity. To produce a model with weather effects but no density-dependent transmission, we eliminated conidia, and instead assumed that resting spores survive for the entire season. Fig. 1 A and B show the predictions of the model with density-dependence only, to show that increases in host density in the density-dependence-only model can strongly increase epizootic intensity. Fig. 1 C, D show the predictions of the model with only weather effects, to show how cool temperatures, high rainfall, and high relative humidity in the weather-only model can similarly increase epizootic intensity (fig. 1 C, D are based on simulated data, but for model-fitting we used observed weather data). Finally, fig. 2 shows the predictions of a model that includes the effects of both density dependence and weather, showing that either increased host density or favorable weather can increase epizootic intensity.

**Fig. 1.**
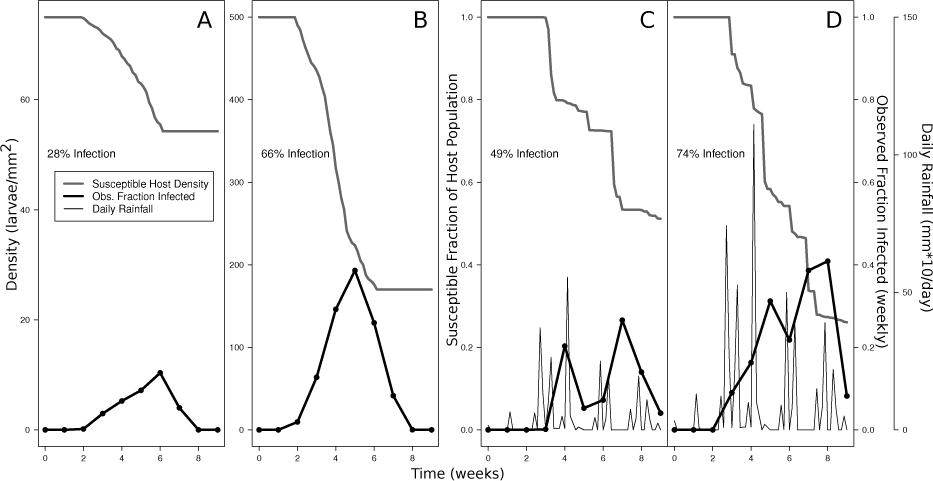
Single realizations of the density-dependence-only model (A, B) and the weather-only model (C and D). In panel A, the initial host density is 75 larvae per square meter, while in panel B the initial host density is 500 larvae per square meter. Panels C and D have the same temporal distribution of rainfall events, but in panel D the volume of rain is twice as high as in panel C.

**Fig. 2.**
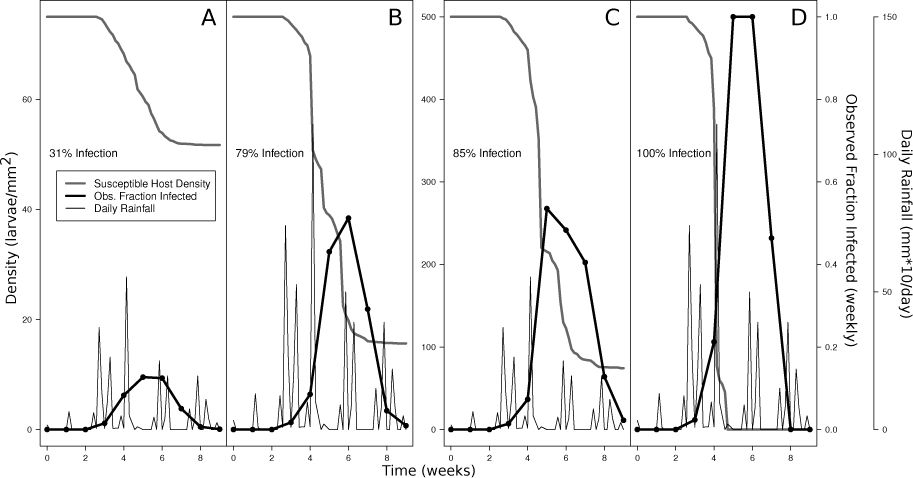
Single realizations of the combined weather and density-dependence model. The four panels demonstrate how the model responds to different combinations of low (A and C) and high (B and D) amounts of rainfall and low (A and B) and high (C and D) initial host densities. Panels A and B have an initial host density of 75 larvae per square meter. Panels C and D have an initial host density of 500 larvae per square meter. In panels B and D, the total amount of rain is twice that in Panels A and C.

Direct inspection of the data confirms that infection rates varied strongly across host densities and weather conditions (fig. 3), which is necessary for our fitting routines to be successful. Also, changes in the force of infection in our experimental data were very close to changes in infection rates in the observational data, suggesting that our experiments provided a reliable estimate of the force of infection. Positive effects of cool, moist weather on transmission are then apparent in comparisons of the rainfall, temperature, and relative humidity data to the infection-rate data (fig. 3), with the proviso that the strong correlations between the three weather variables makes it hard to determine which is the most important.

**Fig. 3.**
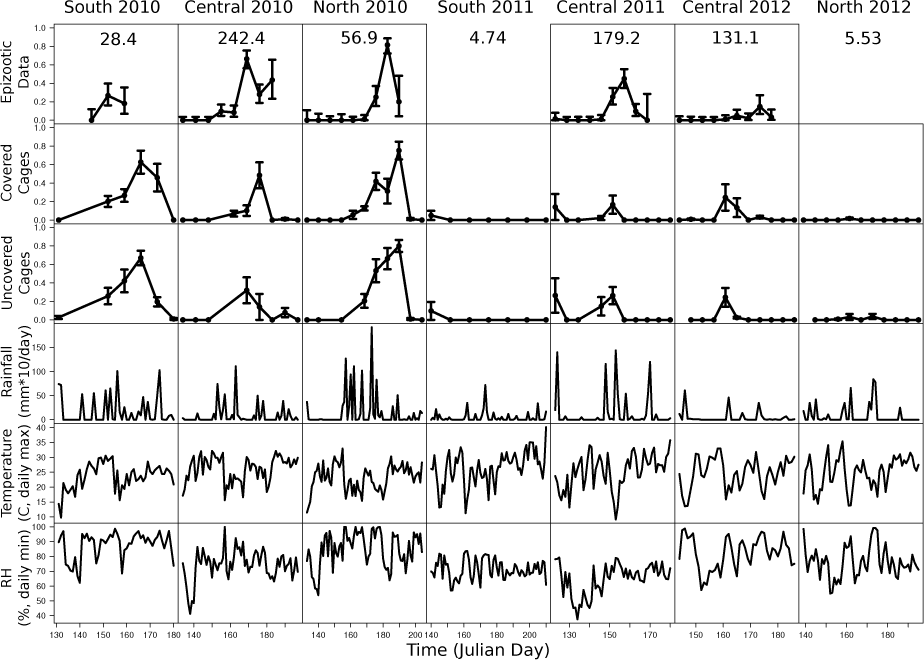
Observational and experimental data on *E. maimaiga* infection rates. Error bars represent one standard error of the mean. Each column shows the estimated initial host density (*S*(0)_*i*_) for each site in the top row in terms of larvae/m^2^. “Covered” refers to an experimental treatment in which plastic buckets were placed over cages to limit exposure to condia, while “Uncovered” cages had no buckets [16]. Bucket effects were modest (see Supplementary Information).

Effects of density, however, are also clearly apparent. First, in the South 2011 and North 2012 sites, larval densities were undetectably low, and apparently as a result the infection rate among experimental larvae was only slightly above zero. This was true even though in both South 2011 and North 2012, the amount of rainfall, the relative humidity and the temperature were similar to what they were in Central 2010, where host densities were high and an intense *E. maimaiga* epizootic occurred. Moreover, in both South 2011 and North 2012, at least a few infections occurred among experimental larvae early in the season, confirming that germinating resting spores were present. The lack of infections among experimental larvae later in the season in both populations emphasizes the importance of host density and conidial transmission in driving epizootics.

Using our data to choose between competing models of *E. maimaiga* transmission first required that we fit the models to the data, which we accomplished using a variant of Metropolis-Hastings Markov-chain Monte Carlo (MCMC) known as “line search-MCMC” [22]. Line search-MCMC uses a large number of line searches to generate automated proposal distributions for MCMC, thereby taking advantage of the highly parallel nature of modern computing environments in a way that is not possible with standard MCMC. To allow for stochasticity, we included daily, stochastic variation in conidial and resting-spore transmission. We then calculated integrated likelihood scores [23] by averaging over realizations, using a stochastic integration routine known as MISER [24] in the gsl scientific computing package [25].

Choosing between models using the AIC model-selection criterion [26] then confirmed that the data are best explained by a model that includes both weather and density-dependent transmission (Table 1). The density-dependence-only model can qualitatively reproduce the data, especially the rise and fall in infection rates over the course of epizootics (fig. 4 “D-D Only”), but it misses the sharp increases in infection rates that result from rainstorms, with their attendant increases in humidity and reductions in temperature. These prediction errors are consistent with the hypothesis that weather affects transmission.

**Table 1.** AIC analysis. The best model is shown in bold-face, while the best weather-only model is shown in italics. The sample size was large enough that the small-sample AIC_c_ criterion gave virtually identical values to standard AIC.

| Model | # Parameters | AIC | $\Delta AIC$ |
| --- | --- | --- | --- |
| Density-dep. only | 23 | 699.8 | 40.6 |
| Weather only |  |  |  |
| Rain | 12 | 737.9 | 78.8 |
| RH | 11 | 706.8 | 47.6 |
| Temp | 11 | 689.4 | 30.3 |
| Rain + RH | 13 | 688.9 | 29.7 |
| *Rain + Temp* | *13* | *678.7* | *19.5* |
| RH + Temp | 13 | 705.4 | 46.2 |
| Rain + RH + Temp | 14 | 682.8 | 23.6 |
| Dens. dep. + Weather |  |  |  |
| Rain | 25 | 701.2 | 42.0 |
| RH | 24 | 662.9 | 3.75 |
| Temp | 24 | 700.5 | 41.3 |
| Rain + RH | 26 | 671.2 | 12.0 |
| Rain + Temp | 26 | 705.7 | 46.5 |
| RH + Temp | 25 | 669.6 | 10.5 |
| **Rain + RH + Temp** | **27** | **659.2** | **0.0** |

**Fig. 4.**
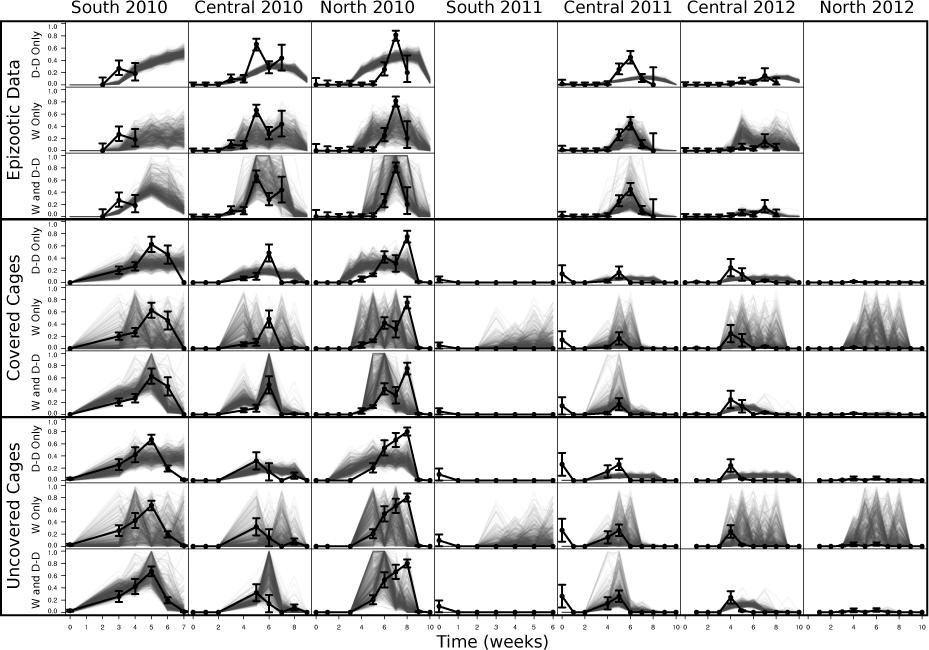
Comparison of multiple realizations of the models to the data. “D-D Only” refers to the density-dependence-only model, “W Only” refers to the best weather-only model, and “W and D-D” refers to the best combined weather and density-dependence model. Black dotted lines are the data (+/-1 SE) and the grey transparent lines are multiple realizations of the models using the best-fit parameter sets.

The best weather-only model in contrast easily reproduces weather-driven increases in infection rates, but it often predicts that infection rates will be high when infection rates in nature were low or zero (fig. 4 “W Only”). Moreover, this overprediction occurs early in the season in study plots with high densities (all plots 2010), over the entire season in plots with low densities (South 2011, North 2012), or late in the season in plots in which densities were apparently not high enough to maintain conidial transmission after the first few weeks (Central 2012). These prediction errors are consistent with the hypothesis that density-dependent transmission and the accumulation of conidia play important roles in transmission.

Although the best combined model is vastly better than either the density-dependence-only model or the best weather-only model, the second-best combined model includes only relative humidity, and it has ΔAIC = 3.75, which indicates that the evidence in favor of the best model, which includes all 3 weather variables, is strong but not overwhelming [26]. Comparisons within the weather-only models are similar, in that the best weather-only model includes rainfall and temperature but not relative humidity, while the weather-only model that also includes humidity has ΔAIC = 4.1 relative to the best weather-only model, showing that the evidence for the best weather-only model is likewise strong but not overwhelming [26]. Notably, however, any model that does not include relative humidity provides a vastly worse explanation for the data than any model that does include relative humidity. Our models thus provide conclusive evidence for an effect of relative humidity, and strong but not conclusive evidence for effects of temperature and rainfall.

An important reason why we were able to detect effects of density on transmission is that density-dependent transmission is required to reproduce the pattern of rising and falling infection rates apparent in the data, emphasizing the usefulness of fitting dynamic, nonlinear models to temporally resolved epizootic data. The inclusion of both resting spores and conidia in our density-dependent models is also important because resting spore transmission rates are almost completely unaffected by the density [14] or even the presence of the insect [28], whereas in previous work density-dependence has only had clear effects during conidia-driven transmission [17, 29].

The effect of the disease-density threshold on the probability of an epizootic, however, is not obvious from the fit of the models to the data, and so we instead plotted model predictions against host density, focusing on cumulative infection rates for consistency with classical theory. As fig. 5 shows, for the best combined density-dependence plus weather model, the qualitative shape of the infection curve is similar to that produced by classical models [6], with the proviso that there is stochastic variation across model realizations that does not occur in classical, deterministic models. As a result, instead of a single threshold value, there is a range of values, below which the probability of a severe epidemic is very low because resting spore transmission is the main source of infection, and above which the probability of a severe epidemic is very high because of repeated rounds of conidial transmission. For this pathogen, weather thus drives variability in the disease-density threshold, but in a more general sense it does not eliminate the threshold effect.

**Fig. 5.**
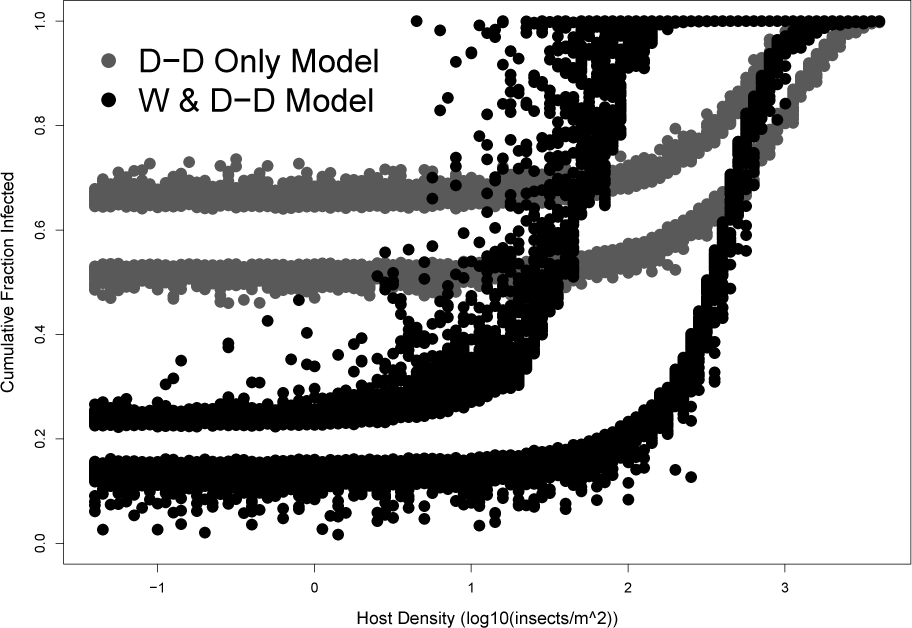
Comparison of the predictions of the density-dependence only model and the best combined model, across a range of densities. Points depict the upper 95% and lower 5% of 1000 realizations at each density. For each simulation of the combined model, we randomly generated daily weather using a Richardson weather generator [27] tuned to the average climate of Michigan. Here we use initial resting spore densities for Central 2010, for which our best estimate for the density-dependence-only model was somewhat lower than for the best combined model (Supplementary Information), emphasizing the importance of resting spore transmission in driving infection at low density in that model.

Lloyd-Smith et al. [7] in contrast argued that demographic stochasticity *is* likely to eliminate the threshold effect, because demographic stochasticity can lead to bimodal distributions of epidemic severity, with a substantial probability of no epidemic even when the host density is well above the threshold. For realistic parameters, however, this bimodality disappears if the initial number infected is more than about 10 individuals, whereas the effects of weather-driven stochasticity are independent of the host population size. It therefore seems likely that weather-driven stochasticity plays at least as important a role in epizootics in nature as demographic stochasticity.

Because our results provide a clear example of how weather stochasticity does not necessarily eliminate threshold effects, we argue that thresholds are of practical use in disease management. For *E. maimaiga* in particular, the existence of a threshold means that it is unlikely that the disease will cause the collapse of rising gypsy moth populations before defoliation occurs. Our best model could nevertheless be used to predict when populations will collapse naturally, reducing control costs, with the proviso that there will be at least modest uncertainty in the extent of the collapse unless accurate weather forecasts are available.

Fig. 5 further shows that, because the density-dependence-only model does not allow for stochastic variation in weather, it produces predictions that are less variable than the predictions of the combined model. The more notable difference between the two models, however, is that, in the density-dependence-only model, infection rates are often high even if host density is low, so that the effect of the disease-density threshold is much weaker than in the combined model. This counter-intuitive effect occurs because the only way that the density-dependence-only model can explain weather-driven variation in infection rates is by allowing for a higher resting-spore transmission rate than in the combined model (Supplementary Information). Fitting density-dependence-only models to data on pathogens that are strongly influenced by weather could thus lead to incorrect inferences about the effects of density.

Before *E. maimaiga* first occurred at high levels in North America in 1989 [30], gypsy moth outbreaks were driven by a combination of a species-specific baculovirus and generalist predators [31]. The high mortality that *E. maimaiga* often causes suggests that it may alter the period or amplitude of outbreaks, but it has only had widespread effects for a few outbreaks, and so the extent to which outbreak dynamics will change is therefore not yet obvious from the data. Theory nevertheless predicts that if *E. maimaiga* transmission were entirely driven by weather, it would serve only as an additional form of stochasticity, leading to greater uncertainty in the timing of outbreaks, but with no consistent change in the mean period or mean amplitude of the population cycle [31]. Our demonstration that *E. maimaiga* transmission is density-dependent in contrast suggests that *E. maimaiga* will likely drive a consistent change in the period and amplitude of gypsy moth outbreaks. Nevertheless, in contrast to previous work [18], our data show that the baculovirus still plays an important role in gypsy moth population dynamics, in that a study population that we had hoped to include in our data set was decimated by the virus, precluding a fungal epizootic. The extent to which *E. maimaiga* will change gypsy moth outbreak cycles is therefore unclear, and so an important next step is to combine our *E. maimaiga* model with models of the gypsy moth baculovirus [31].

## Methods

### Model Structure

Our initial efforts to fit mechanistic models to our data made clear that the sharp increase in infection rate over time in most plots and years could not be easily captured unless we followed Weseloh [32] in assuming that the conidial transmission rate *v_C_* increased with time, reflecting increases in larval size that make it easier for conidia to contact later instars. We therefore assumed that *ν_C_* increases with dd10, the number of accumulated degree days over 10° C, the lowest temperature at which infections occur, scaled by the size of the fourth instar (i.e., stage) larvae that we used in our experiments.

We then added stochasticity to our models by multiplying the resting spore and conidial transmission rates *ν_R_,_t_* and *ν_C_,_t_* by a stochasticity term, *e^ϵt^*. Here *ϵ_t_* is a normally distributed random variate with mean zero and standard deviation σ, so that *e^ϵ_t_^* varies between 0 and ∞, and *σ* is fit to the data. By drawing new values of *ϵ;_R_,_t_* and *ϵ_C_,_t_* every day, the model generates a new, random value for each daily transmission rate.

The simplest version of our transmission-rate functions occurs in the density-dependence only model:

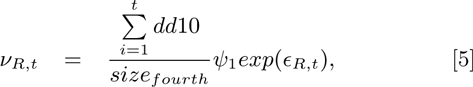

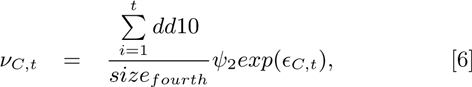

The coefficients *ψ*_1_ and *ψ*_2_ describe the rate at which transmission rates increase with increases in larval size, while *size_fourth_* is the average weight of a fourth instar larva.

In the weather-only models, we eliminated density-dependence in transmission by assuming that the force of infection does not depend on the density of infectious spores, whether resting spores or conidia. Because in this model we have effectively eliminated the distinction between conidia and resting spores, we instead symbolize transmission as *ν_F_,_t_*, where *F* stands for “fungus”:

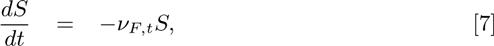

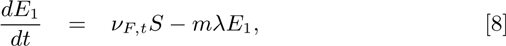

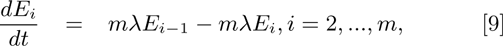

To make transmission *ν_F_,_t_* a function of weather, we defined the following functions, which translate our three weather variables, precipitation, relative humidity (RH), and temperature, into transmission rates:

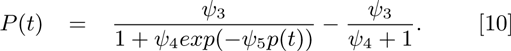

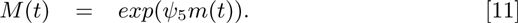

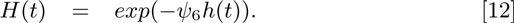

Here *m*(*t*) and *h*(*t*) represent daily values of minimum RH and maximum temperature, respectively, while *p*(*t*) represents cumulative rainfall over the preceding 10 days, based on work of Reilly et al. [16], and the coefficients *ψ*_3_-*ψ*_6_ are fit parameters. Using the minimum RH and the maximum temperature ensured high variability in weather effects between our sites, thereby increasing our chances of accurately quantifying the effects of weather on transmission. Following previous work, we assumed that transmission increases logistically with increases in accumulated daily rainfall over the preceding 10 days [16, 28], with the proviso that *P*(*t*) is 0 when no rain has fallen (*p*(*t*) = 0). Also, because Hajek et al. [33] found that conidia production is positively correlated with humidity, we assumed that conidial transmission increases exponentially with increasing RH. Finally, high temperatures have been found to inhibit conidia production [33], and so we made conidial decay an exponentially declining function of daily maximum temperature.

We then included enough models so that *ν_F_,_t_* was a product of all possible combinations of the weather-variable functions. For example, in the model containing all weather variables, we have:

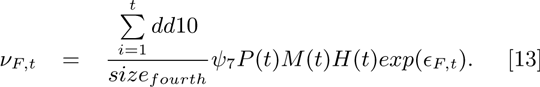

Here we again allowed for stochasticity, such that *ϵ_F,t_* is a normally distributed random variate with mean 0 and standard deviation *σ_F_*.

To allow weather to affect transmission in our density-dependence plus weather models, we used similar functions to those in the weather-only models, except that in the density-dependence plus weather models we again distinguished between resting spores and conidia. We therefore have different transmission functions for resting spores and conidia, such that resting spore germination is an increasing function of rainfall, while conidial infection rates depend on humidity, and conidial survival rates depend on temperature:

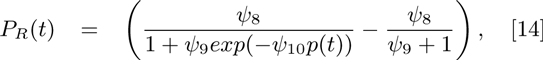

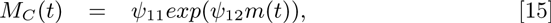

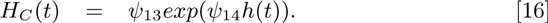

Here *P_R_*(*t*) is the dependence of resting spore transmission on rainfall, *M_C_*(*t*) is the dependence of conidial transmission on humidity, and HC(t) is the dependence of conidial survival on temperature. The coefficients ψ_8_-ψ_14_ are then fit parameters.
Because we again include the effects of increasing larval size on larval susceptibility, for this model the transmission and survival parameters become:

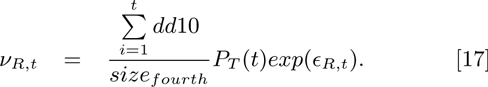

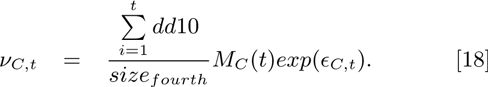

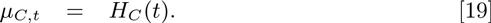

We calculated starting densities of resting spores *R*(0) by treating the initial density as a site-specific fit parameter. Resting spores in nature are only active for a few weeks each spring, and so for simplicity we assumed that the resting spores were germinating in each plot only over a fixed period, defined by the parameters *T_R,start_* and *T_R,end_* (Supplementary Information).

### Data Collection

Larvae collected in naturally occurring populations were reared individually on artificial diet in cups with tightly fitting lids at 21°C in the laboratory. Larvae that die and produce conidia are visually distinct, but to ensure accurate identification of cause of death, we examined smears from dead larvae at 400× under a light microscope to look for conidia or resting spores. In the experimental part of our study, we standardized infection risk across weeks by using only fourth instars, and by using larvae from egg masses collected each season from the same field population of gypsy moths in Roscommon County, Michigan. Because larvae in natural populations spanned a range of instars over time, we allowed for differences in infection risk between feral and experimental larvae in our models by including additional parameters that described the transmission rate in each experimental treatment.

### Model-Fitting Algorithm

To fit our models to data, we used maximum likelihood, such that each likelihood was based on both experimental and observational data. In calculating likelihoods, our goal was to integrate over the stochasticity, thereby calculating an integrated likelihood [23], according to:

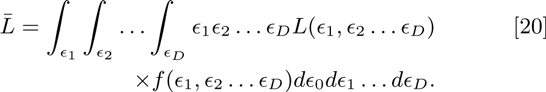

Here the average likelihood 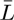 is calculated from the likelihood in each realization 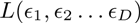, which in turn depends on *ϵ_i_*, the value of the stochasticity in day *i* of each realized epizootic, so that 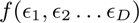 is the (Gaussian) frequency distribution of the *ϵ_i_’S*. Evaluation of the integral in equation (21) using numerical integration methods, however, is impractical [24], because of the complexity of our models. We therefore instead used the Monte Carlo MISER integration algorithm [25], which is based on recursive, stratified sampling [24]. To find the maximum likelihood of each model, we then used line-search MCMC, as described in the main text [22]. Inspection of trace plots and calculation of Gelman-Rubin statistics (all ≤ 1.04) confirmed that our routines converged.

### Simulating Weather

To use our models to understand the effects of variability in weather conditions on model predictions, we generated artificial weather using a Richardson weather generator. To tune this weather generator so that it produced realistic weather data for our study areas [27], we set the daily probability of rainfall during the larval period to be 40%, the frequency of rainfall that we observed in our weather stations. If a rainfall event occurred, we drew the amount of rain that fell (*p*(*t*)) from a log-normal distribution with a mean and variance that we varied to understand the effects of rainfall intensity. We then estimated daily temperature (*h_ave_*(*t*), *h_max_*(*t*)) and RH (*m*(*t*)) values from the generated rain data using correlations between rain and the other weather variables estimated by fitting linear models to our observed weather data (Supplementary Information).

## Acknowledgments.

Our work was supported by a National Science Foundation Graduate Student Research Fellowship to C.K., and by NIH grant #R01GM96655 awarded to G. Dwyer, V. Dukic and B. Rehill. Crucial assistance was provided by the Michigan Department of Natural Resources.

## Supplementary Information

**Model Parameter Values**. Tables 2-4 show the best-fit parameter values for the density-dependence only model, the weather-only model, and the combined density-dependence plus weather model.

**Table 2.** Parameter Values for Density-Dependence Only Model

| Parameter |  | Median | 95% HPD |
| --- | --- | --- | --- |
| $\nu_{R,t}$ | $\psi_1$ | 12.9 | [11.7, 14.2] |
| | $\epsilon_{R,t}$ | 0.494 | [0.489, 0.499] |
| $\nu_{C,t}$ | $\psi_2$ | $1.96 \times 10^{-3}$ | $[1.76 \times 10^{-3}, 2.17 \times 10^{-3}]$ |
| | $\epsilon_{C,t}$ | 0.205 | [0.162, 0.259] |
| $\mu_{C,t}$ | | 0.404 | [0.314, 0.510] |
| $\lambda$ | | 0.109 | [0.0974, 0.140] |
| $size_{fourth}$ | | 301.3 | [292.1, 311.1] |
| $T_{R,start}$ | | 120.4 | [118.8, 122.1] |
| $T_{R,end}$ | | 472.9 | [470.3, 475.7] |
| $T_{C,end}$ | | 336.2 | [333.4, 338.9] |
| $\gamma_1$ | | 1.29 | [1.17, 1.40] |
| $\gamma_2$ | | 0.706 | [0.657, 0.764] |
| $\theta_1$ | | 2.12 | [1.59, 2.87] |
| $\theta_2$ | | 7.65 | [6.55, 8.89] |
| $\theta_3$ | | 3.71 | [3.19, 4.29] |
| $\theta_4$ | | 6.10 | [5.46, 6.88] |
| $R(0)$ | 2010 South | $4.26 \times 10^{-3}$ | $[3.70 \times 10^{-3}, 4.92 \times 10^{-3}]$ |
| | 2010 Central | $1.52 \times 10^{-3}$ | $[1.44 \times 10^{-3}, 1.61 \times 10^{-3}]$ |
| | 2010 North | $3.75 \times 10^{-3}$ | $[3.52 \times 10^{-3}, 4.00 \times 10^{-3}]$ |
| | 2011 South | $1.08 \times 10^{-12}$ | $[1.05 \times 10^{-12}, 1.12 \times 10^{-12}]$ |
| | 2011 Central | $5.54 \times 10^{-4}$ | $[5.01 \times 10^{-4}, 6.07 \times 10^{-4}]$ |
| | 2012 Central | $6.33 \times 10^{-4}$ | $[5.97 \times 10^{-4}, 6.70 \times 10^{-4}]$ |
| | 2012 North | $1.15 \times 10^{-4}$ | $[9.18 \times 10^{-5}, 1.43 \times 10^{-4}]$ |

**Table 3.** Parameter values for the best weather-only model (Rain + Temp)

| Parameter |  | Median | 95% HPD |
| --- | --- | --- | --- |
| $\nu_{F,t}$ | $\psi_3$ | 1.60 | [1.36, 1.88] |
| | $\psi_4$ | 2161.4 | [535.5, 8481.6] |
| | $\psi_6$ | 0.138 | [0.111, 0.171] |
| | $\psi_7$ | 0.247 | [0.233, 0.263] |
| | $\epsilon_{F,t}$ | 0.543 | [0.532, 0.556] |
| $\lambda$ | | 0.102 | [0.0896, 0.116] |
| $size_{fourth}$ | | 280.2 | [270.9, 289.3] |
| $T_{R,start}$ | | 107.2 | [104.1, 110.3] |
| $T_{R,end}$ | | 463.5 | [456.9, 472.7] |
| $\gamma_1$ | | 1.14 | [0.775, 1.67] |
| $\gamma_2$ | | 0.854 | [0.796, 0.917] |
| $\theta_1$ | | 10.2 | [8.67, 12.0] |
| $\theta_3$ | | 8.48 | [7.62, 9.48] |

**Table 4.** Parameter values for combined weather plus density-dependence model (Rain + RH + Temp)

| Parameter | Median | 95% HPD |
| --- | --- | --- |
| $\nu_{R,t}$ | | |
| $\psi_8$ | 3.71 | [2.64, 5.00] |
| $\psi_9$ | 3.44 | [1.46, 7.87] |
| $\psi_{10}$ | 0.176 | [0.132, 0.223] |
| $\epsilon_{R,t}$ | 0.372 | [0.260, 0.519] |
| $\nu_{C,t}$ | | |
| $\psi_{11}$ | $2.40 \times 10^{-4}$ | $[1.76 \times 10^{-4}, 3.25 \times 10^{-4}]$ |
| $\psi_{12}$ | $7.01 \times 10^{-2}$ | $[6.50 \times 10^{-2}, 7.57 \times 10^{-2}]$ |
| $\epsilon_{C,t}$ | 0.192 | [0.177, 0.208] |
| $\mu_{C,t}$ | | |
| $\psi_{13}$ | $9.47 \times 10^{-3}$ | $[5.60 \times 10^{-3}, 1.57 \times 10^{-2}]$ |
| $\psi_{14}$ | 0.235 | [0.216, 0.256] |
| $\lambda$ | 0.122 | [0.110, 0.135] |
| $size_{fourth}$ | 291.1 | [281.942, 301.2] |
| $T_{R,start}$ | 100.1 | [94.5, 105.8] |
| $T_{R,end}$ | 267.3 | [261.2, 273.8] |
| $T_{C,end}$ | 524.5 | [514.7, 535.5] |
| $\gamma_1$ | 2.23 | [2.01, 2.46] |
| $\gamma_2$ | 0.943 | [0.896, 0.995] |
| $\theta_1$ | 8.33 | [7.53, 9.20] |
| $\theta_2$ | 7.97 | [6.95, 9.01] |
| $\theta_3$ | 7.01 | [6.39, 7.82] |
| $\theta_4$ | 6.48 | [5.59, 7.58] |
| $R(0)$ | | |
| 2010 South | $1.91 \times 10^{-2}$ | $[1.52 \times 10^{-2}, 2.36 \times 10^{-2}]$ |
| 2010 Central | $5.69 \times 10^{-3}$ | $[4.43 \times 10^{-3}, 7.36 \times 10^{-3}]$ |
| 2010 North | $1.87 \times 10^{-3}$ | $[1.53 \times 10^{-3}, 2.29 \times 10^{-3}]$ |
| 2011 South | $1.09 \times 10^{-12}$ | $[9.83 \times 10^{-13}, 1.20 \times 10^{-12}]$ |
| 2011 Central | $2.26 \times 10^{-3}$ | $[1.90 \times 10^{-3}, 2.70 \times 10^{-3}]$ |
| 2012 Central | $4.41 \times 10^{-3}$ | $[3.81 \times 10^{-3}, 5.14 \times 10^{-3}]$ |
| 2012 North | $7.00 \times 10^{-4}$ | $[4.62 \times 10^{-4}, 1.09 \times 10^{-3}]$ |

### Plot locations

In each year, we attempted to find a southern population, a central population, and a northern population, across what was effectively a 200km transect. Because some populations collapsed each year, in practice, this meant that we searched within the southern, central and northern parts of the lower peninsula of the state of Michigan, USA to find a population in each general area, a search that was unsuccessful in the southern part of the peninsula in 2012. Plot locations are shown in fig. 6, and the latitudes, longitudes and distances between plots are given in Table 5

**Table 5.** Latitudes (“Lat.”) and longitudes (“Long.”) between our study plots, together with distances from the nearest plot to the south (“Dist from S/C”) and the total south to north distance (“Total S/N Dist.”)

| Year | Location | Lat. | Long. | Dist. from S/C | Total S/N Dist. |
| --- | --- | --- | --- | --- | --- |
| 2010 | South | 42.36 | -85.35 |  |  |
|  | Central | 44.46 | -84.60 | 240.96 |  |
|  | North | 45.48 | -84.68 | 113.57 | 354.53 |
| 2011 | South | 42.61 | -85.45 |  |  |
| 2011 | Central | 44.47 | -84.60 | 217.04 |  |
| 2011 | North | 45.19 | -84.23 | 85.44 | 302.48 |
| 2012 | Central | 44.47 | -84.60 |  |  |
| 2012 | North | 45.48 | -84.68 | 85.44 | 85.44 |

**Fig. 6.**
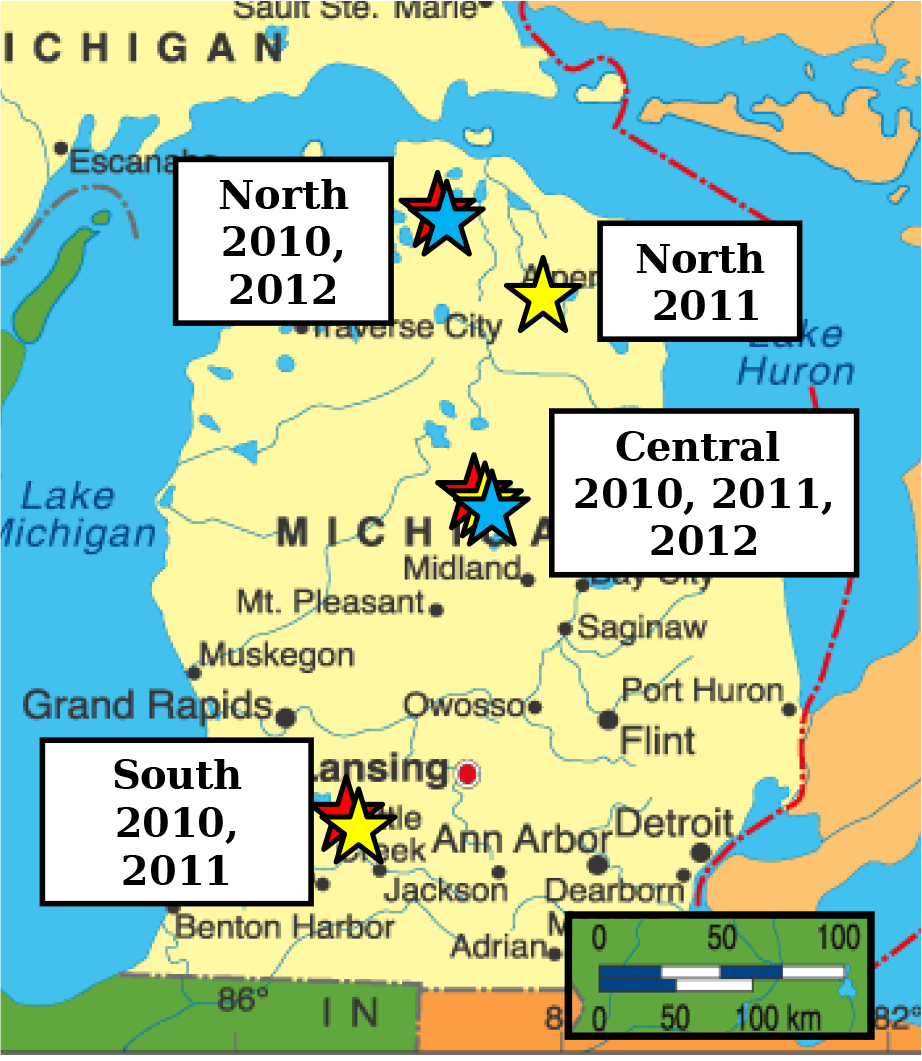
Plot locations.

### Data Collection

To ensure that we were able to collect 100 larvae each week at each site, we first carried out qualitative surveys in the winter before each larval season, to roughly assess local densities. At each site, we then selected a square plot 500m × 500m in area within which all larvae would be collected during the larval period. To quantify initial larval densities within this area, we carried out egg mass counts on five 1/40th hectare circles within each square, locating one circle in each corner and one near the center of the square. Previous work has shown that egg mass counts within 1/40th hectare plots provide reliable predictors of larval densities [13]. The average of these counts served as our estimates of the initial density S(0), in terms of egg masses per square meter, at each each site. To convert to units of larvae per square meter, we assumed that 400 larvae hatch from each egg mass [13], but this assumption affects only our parameter values and not our conclusions [6]. To monitor weather variables, we placed a weather station at the center of the square at each site, and we recorded rainfall, temperature and relative humidity. The station consisted of a RainWise tipping-bucket rain gauge connected to a RainWise RainLog data logger and a HOBO U23 Pro v2 External Temperature/Relative Humidity Data Logger - U23-002 which logged conditions every 5 minutes.

To quantify natural *E. maimaiga* infection rates, we sampled 100 larvae from the population at each site in each week. We found larvae by searching the ground, foliage and tree trunks and we captured them in individual 2 oz plastic cups containing artificial agar-based gypsy moth diet [34]. Larvae were then reared in the laboratory in these cups until death or pupation at 21° C, a temperature that maximizes the chance that an infection will successfully produce conidia [14]. Larvae were checked for death and signs of conidia twice a week for three weeks. Conidia are often visually apparent, but even when they are not, they can be easily seen at 400× under a light microscope, as can resting spores [14].

Preliminary work showed that estimating the conidial transmission rate *ν_C_* and decay rate *μ_C_* from observational data alone leads to high uncertainty in each parameter, suggesting that, when it comes to observational data, the two parameters are close to being nonidentifiable. This is important, because transmission and decay may have very different dependence on climate variables, and so nonidentifiability could lead to misidentification of the effects of weather. We there-fore collected experimental data from each plot, by exposing lab-reared larvae to the environment for 24 hours. This period is short enough relative to the speed of kill of the disease that the resulting data provide a close approximation of the force of infection of the disease.

Larvae for experiments were hatched from eggs collected from a population in Roscommon County, Michigan, 10 kilometers from our study site there. To remove any baculovirus from the eggs, we disinfected them in a 4% by volume formaldehyde solution [35]. Hatching larvae were then reared on artificial diet in the laboratory at 25° C in a different room from field-exposed insects, which ensured that the lab-reared insects did not contact either the fungal or viral pathogens until they were deployed.

Cages were made of aluminum screen, 20 × 20 × 5 cm, containing approximately 20 unexposed lab-reared larvae, and were placed on the soil at the base of a tree each week. To control for differences in the susceptibility of different-aged larvae, we used only recently molted fourth instars. Because red oak (*Quercus rubra*) is a most preferred gypsy moth host tree species [36], trees were chosen to be the overstory red oak within each of the five egg-mass-count circles at each site.

We placed two cages under each tree each week. Following Reilly et al. [16], one cage per tree was covered with a clear plastic box to reduce exposure of larvae to air-borne conidia while the other was uncovered so that larvae were exposed to both conidia and resting spores. After 24 hours, larvae were removed to the laboratory, where they were reared separately on artificial diet at 21° C until pupation or death, in the same manner as larvae collected from feral populations at each site. To ensure that we were able to separately estimate conidial transmission and conidial decay, we deployed experimental larvae from the time of larval hatch until two weeks after naturally occurring larvae had pupated at each site.

### Model Details

As we described in the main text, resting spores are only active for a few weeks during the larval period, but the mechanisms determining the beginning and end of this period are unknown [14]. In our field collections, however, we observed that the time of initiation (*T_R,start_*) and cessation (*T_R,end_*) of resting spore infection was positively correlated with latitude. Accordingly, because gypsy moth hatch time and development depend on the number of accumulated growing degree days above 10° C (dd10) [13], which is in turn a function of latitude, we set the resting spore activity in our models to begin and end after an estimated value of dd10 had been reached.

Again as we described in the main text, infections in fifth and sixth instar larvae do not produce infectious conidia, instead producing resting spores for the next season [21]. To incorporate this behavior, we fit a parameter to estimate the dd10 required for our host populations to reach the fifth instar (*T_c,end_*), after which cadavers cease producing new infectious conidia, although existing conidia may continue to cause infections.

### Likelihood Function

In theory, a binomial distribution should be an appropriate likelihood function, because the pathogen must kill its host to be transmitted. The binomial distribution, however, assumes individual hosts are independent, which may be untrue if, for example, individuals that have either higher or lower infection risk than average cluster together within a population [37]. This lack of independence can cause the true variance in infection risk to be substantially higher than the variance assumed under the binomial, a phenomenon kinown as over-dispersion [38]. In the absence of direct information on over-dispersion, a useful approach is to use a beta-binomial distribution, which is derived from the binomial by assuming that p, the probability of an infection in the binomial, follows a beta distribution in which the quantity in question varies between 0 and 1. The resulting distribution has two parameters, as opposed to the single parameter of the binomial.

The additional parameter allows us to increase the variance as needed to explain the measurement error in the data, which would be reflected in a lack of fit of a model to the data. To explain this approach, we define the parameters of the beta-binomail as *a* and *b*. We then re-parameterized according to:

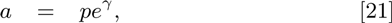

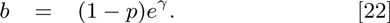

Here *p* is again the probability of infection in the binomial, while *γ* is a parameter that affects the variance of *p*. Under this re-parameterization, if the sample size is *n*, then under both the binomial and the beta-binomial, the expected number responding is *np*. The variance of the binomial, however is *np*(1−*p*), whereas the variance of the beta binomial is instead:

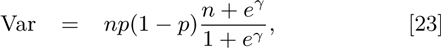

This expression makes clear that, as 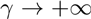, the variance of the beta binomial approaches the variance of the binomial, but as 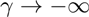, the variance instead approaches *n*^2^*p*(1−*p*). By adjusting *γ*, we can thus adjust the variance of the beta-binomial as needed to explain the lack of fit between the model and the data in any realization. Because the models attempt to predict the value of *p*, *γ* is thus an inverse measure of the variance of the measurement process, and therefore quantifies the the measurement error. Because we do not have independent information about the value of *γ*, we fit *γ* to the data, along with the other parameters.

### Including Observational and Experimental Data

Because we have both observational and experimental data, we calculated two likelihood scores for each parameter set, one for the observational data and one for the experimental data. Given the model, the two data sets are independent and could therefore be summed on a log scale. In practice, however, we expected that conditions in our experiments would be at least slightly different from conditions in nature, and so we constructed separate models for the experimental insects.

Because the experimental insects were only deployed in the field for 24 hours, none of them died of the disease, and so they did not contribute to the conidia population. Meanwhile, our data consist only of the fraction infected, and so the length of time that the experimental insects spend in the exposed categories is of no interest. We nevertheless expected that the infection rates would be at least moderately lower in the covered cages, and so we included an equation for both covered and uncovered cages:

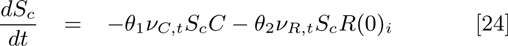

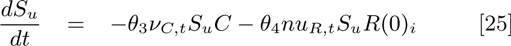

Here *S_c_* is the density of insects in the covered cages, and *S_u_* is the density of the insects in the uncovered cages. As in the main text, *R*(0)_*i*_ is the density of resting spores in plot i, and *C* is the density of conidia, where changes in *C* are calculated using equations (1)-(4) in the main text. The parameters *θ*_1_ and *θ*_2_ describe the change in the conidial transmission rate *ν_C_,_t_* and the resting spore transmission rate *ν_R,t_* in the covered cages relative to the corresponding transmission rates in nature, while the parameters *θ*_3_ and *θ*_4_ describe the changes in the uncovered cages.

### Linear models for weather variables

To account for autocorrelation and covariate-correlation among weather variables when building our weather generator, we compared the ability of different time-series models of varying complexity to explain the data logged by our weather stations at each field site [26]. This approach showed that relative humidity and temperature are best predicted from rainfall. In our simulations, we therefore used a log-normal to describe rain *R*(*t*) on day *t*, with a mean of 25.87 mm per day, and with variance 1189.2. We then generated values of maximum relative humidity *RH_max_*(*t*), average temperature *T*_avg_(*t*) and maximum temperature *T*(*t*) on day *t* from the following regressions based on our data:

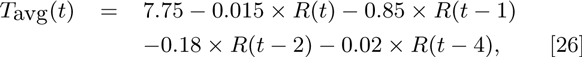

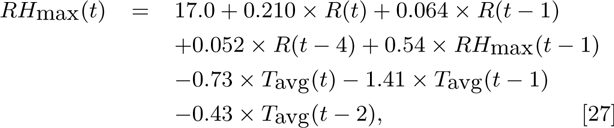

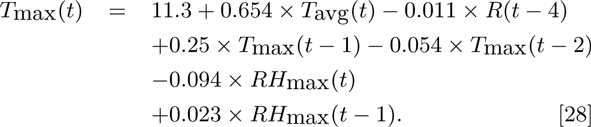

